# A common neural network among state, trait, and pathological anxiety from whole-brain functional connectivity

**DOI:** 10.1101/158055

**Authors:** Yu Takagi, Yuki Sakai, Yoshinari Abe, Seiji Nishida, Ben J. Harrison, Ignacio Martínez-Zalacaín, Carles Soriano-Mas, Jin Narumoto, Saori C. Tanaka

**Author notes:** Corresponding author (Yu Takagi), (Saori C. Tanaka).

## Abstract

Anxiety is one of the most common mental states of humans. Although it drives us to avoid frightening situations and to achieve our goals, it may also impose significant suffering and burden if it becomes extreme. Because we experience anxiety in a variety of forms, previous studies investigated neural substrates of anxiety in a variety of ways. These studies revealed that individuals with high state, trait, or pathological anxiety showed altered neural substrates. However, no studies have directly investigated whether the different dimensions of anxiety share a common neural substrate, despite its theoretical and practical importance. Here, we investigated a neural network of anxiety shared by different dimensions of anxiety in a unified analytical framework using functional magnetic resonance imaging (fMRI). We analyzed different datasets in a single scale, which was defined by an anxiety-related neural network derived from whole brain. Through the fMRI task for provoking anxiety, we found a common neural network of state anxiety across participants (1,638 trials obtained from 10 participants). Then, using the resting-state fMRI in combination with the participants’ trait anxiety scale (879 participants from the Human Connectome Project), we demonstrated that trait anxiety also shared the same neural network as state anxiety. Furthermore, the common neural network between state and trait anxiety could detect patients with obsessive-compulsive disorder, which is characterized by pathological anxiety-driven behaviors (174 participants from multi-site datasets). Our findings provide direct evidence that different dimensions of anxiety are not completely independent but have a substantial biological inter-relationship. Our results also provide a biologically defined dimension of anxiety, which may promote further investigation of various human characteristics, including psychiatric disorders, from the perspective of anxiety.

## 1. Introduction

Anxiety is a future-oriented mental state activated by distant and potential threats rather than specific and predictable ones (Calhoon and Tye, 2015). On the one hand, anxiety drives us to avoid frightening situations and to achieve our goals. On the other hand, excessive anxiety may cause distress and impairment in daily life. Although anxiety is a common mental state in humans, we experience anxiety in a variety of forms. For example, public speaking or leaving one’s home can induce anxiety related to a fear of negative evaluation by others and risk of theft, respectively. Such anxieties may drive us to do something to overcome our anxiety, such as practicing a speech or repeatedly checking that the house door is locked.

One conventional way to study anxiety is to investigate the state associated with the feeling of anxiety, namely, state anxiety. Other studies focus on the frequency of anxiousness, namely, trait anxiety, which is measured using self-report questionnaires. Another major research field concerns the anxiety of patients with psychiatric disorder. This type of anxiety includes social anxiety disorder, obsessive-compulsive disorder (OCD), and generalized anxiety disorder. Such “pathological anxiety”, which is defined by clinically significant levels of anxiety (i.e., excessive, uncontrollable anxiety), incurs tremendous socioeconomic costs (Greenberg et al., 1999) and roughly 30% of people experience an anxiety-related disorder at some point in their lifetime (Kessler et al., 2005).

In the neuroscience field, using functional magnetic resonance imaging (fMRI), different dimensions of anxiety have been investigated by several types of approaches, which correspond to the different dimensions of anxiety described above. To investigate the brain activity underlying state anxiety, researchers have experimentally induced participant’s anxiety inside of the MRI scanner (Mataix-Cols et al., 2003; Satpute et al., 2012). Other studies focused on the relationship between trait anxiety and brain activity (Baur et al., 2013; Kim et al., 2011; Modi et al., 2015; Tian et al., 2016; Yin et al., 2016). These latter studies have recently focused on changes in brain activity, measured by resting-state fMRI (rs-fMRI). Finally, other studies have focused on alterations in the neural substrate in populations with pathological anxiety and have revealed differences in neural activity compared with healthy populations (Banca et al., 2015; Beucke et al., 2013; Cha et al., 2014; Etkin et al., 2010; Giménez et al., 2012; Hahn et al., 2011; Harrison et al., 2009; Liu et al., 2015; Sakai et al., 2011; Wang et al., 2016).

Previous findings have suggested an interaction among state, trait, and pathological anxiety (e.g., Mathews, 1990; Williams et al., 1996). If there is a common biological substrate, it has the potential to be used for risk assessment and early detection of pathological anxiety, which would enable its prevention before onset or its evaluation for treatment. From a theoretical perspective, it would also be helpful to understand the dynamics of the development of pathological anxiety. In addition, given that the hypothesis of a psychiatric disorder spectrum is gaining attention (Adam, 2013), such a biologically defined index would provide an objective, reliable biological dimension of anxiety for the spectrum, which may be valuable for understanding various human characteristics, including psychiatric disorders. However, no studies have directly investigated whether there is a common biological substrate among state, trait, and pathological anxiety.

To determine whether there is a common biological substrate among state, trait, and pathological anxiety, we adopted a single neuronal index defined by anxiety-related functional connectivity (FC), a measure of temporally correlated fluctuations in blood oxygen level-dependent (BOLD) signal among different regions. FC has been successfully used to elucidate neural mechanisms of various individual characteristics (Cole et al., 2012; Finn et al., 2015; Liem et al., 2016; Rosenberg et al., 2016). Indeed, FCs have been correlated with individual differences in state or trait anxiety (Baur et al., 2013; Modi et al., 2015; Satpute et al., 2012; Tian et al., 2016) and are different between healthy people and individuals with pathological anxiety (Banca et al., 2015; Beucke et al., 2013; Cha et al., 2014; Etkin et al., 2010; Giménez et al., 2012; Hahn et al., 2011; Harrison et al., 2009; Liao et al., 2010; Liu et al., 2015; Sakai et al., 2011; Wang et al., 2016).

Here, in a data-driven manner, we directly tested the hypothesis that there is a common neural network among state, trait, and pathological anxiety. We first investigated whether state and trait anxiety shared a common neural network in healthy people. Then, to investigate how state and trait anxiety are involved in pathological anxiety, we tested which sets of FCs were generalized to patients with OCD, a disorder characterized by pathological anxiety-driven behavior, among the neural networks related to state and/or trait anxiety.

## 2. Materials and methods

### 2.1. Anxiety provocation task

#### Participant recruitment

To effectively extract anxiety-related neural networks, we recruited 10 healthy participants (9 men, ages 20–24 years, mean age 22.2 years) who tended to be anxious in their daily life from 432 volunteers who completed a questionnaire prior to the fMRI experiments. Specifically, we recruited participants who had a score of greater than or equal to 80 on the Padua Inventory (Burns et al., 1996) or 13 on the Maudsley Obsessional Compulsive Inventory (Hodgson and Rachman, 1977). All participants were primarily evaluated using the Structured Clinical Interview for DSM-IV Axis I Disorders-Non-Patient Edition (SCID-NP) (First et al., 2002). No participant had a current DSM-IV Axis I diagnosis of any significant psychiatric disorder. Participant consent was obtained in accordance with a protocol reviewed and approved by the Ethics Committee of the Advanced Telecommunications Research Institute International. At the time of the experiments, the mean ± standard deviation of the Padua Inventory and Maudsley Obsessional Compulsive Inventory were 71.5 ± 23.6 and 13.2 ± 2.3, respectively.

#### 2.1.1. Stimuli selection

Previous studies demonstrated that neural activity related to anxiety can be extracted using fMRI by presenting two sets of stimuli to participants (Blair et al., 2008; Giménez et al., 2012; Lorberbaum et al., 2004; Mataix-Cols et al., 2003; Rotge et al., 2008; Tillfors et al., 2001). One set of stimuli is for provoking anxiety (anxiety stimuli), for example, the image of a key hole, which may induce anxiety about theft. The other set of stimuli is a relatively neutral stimulus, for example, the image of a nature scene. Here, we conducted a similar task, by iteratively presenting two sets of stimuli to the participants while controlling the semantics (categories of the objects) and basic features (color and luminance) of the two sets of stimuli (Fig. 1a; see Supplementary Notes). Before the fMRI experiments, all participants rated their subjective anxiety in response to approximately 200 images in a Likert scale between 1 and 9 (1, lowest anxiety; 9, highest anxiety). Then, for each participant, anxiety images, those with the top 50 anxiety scores, and neutral images, those with the bottom 50 anxiety scores, were selected as stimuli for the fMRI.

**Figure 1.**
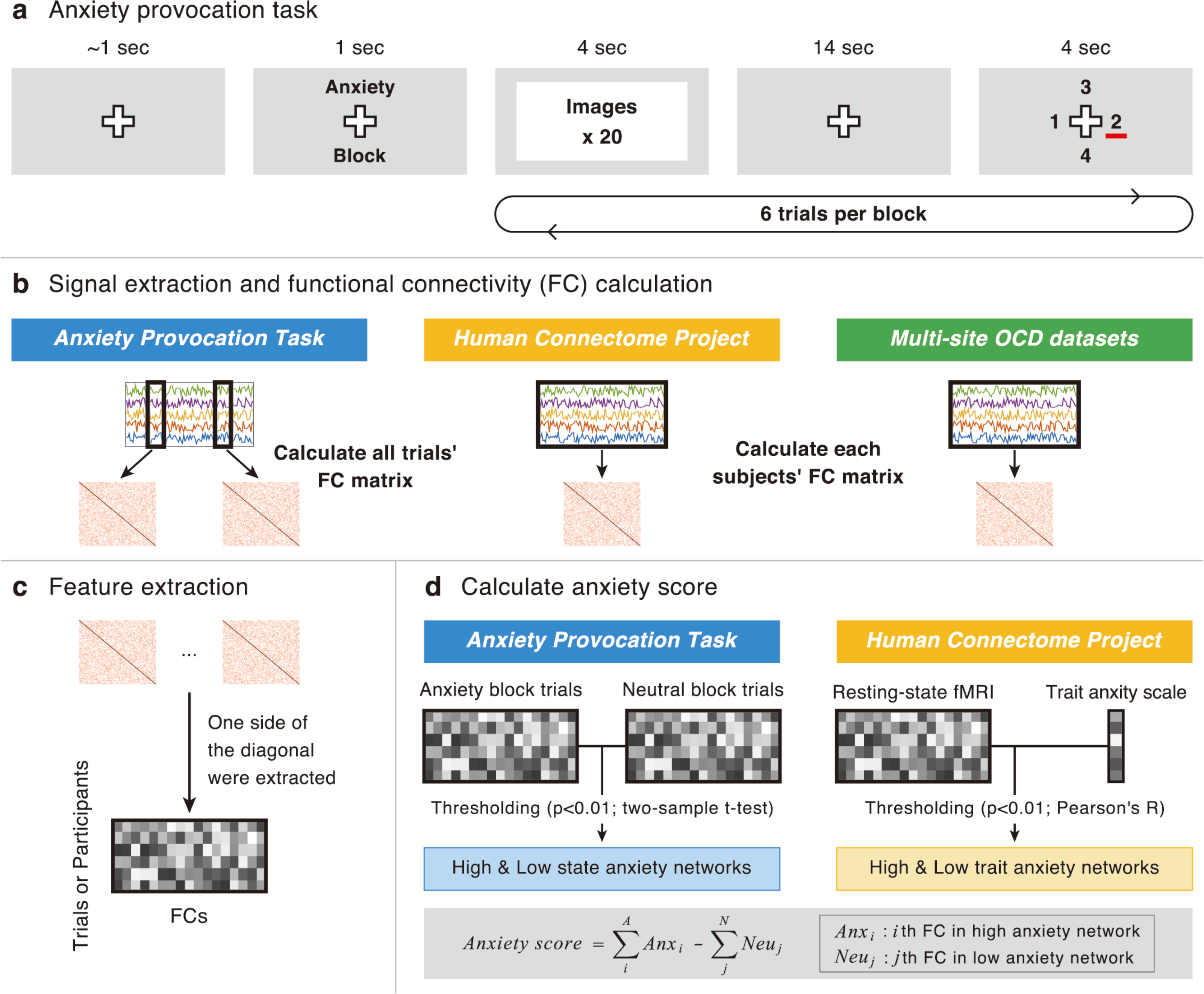
Experimental paradigm and definition of anxiety score. **(a)** The sequence of events for a given trial of the anxiety provocation task. **(b–d)** Schematic diagram of the analyses. **(b)** Three types of datasets were used, that is, anxiety provocation task, Human Connectome Project, and multi-site OCD datasets. Signals of interest were extracted from all participants. The signals were then turned into functional connectivity (FC) via covariance estimation. Note that trial-wise FC matrices were estimated for the anxiety provocation task. In contrast, participant-wise FC matrices were estimated for the Human Connectome Project dataset and multi-site OCD datasets. **(c)** FC matrices were turned into the feature matrix for the subsequent analysis. **(d)** FCs with a strong relationship to anxiety were selected via statistical thresholding through cross-validation. Anxiety scores for trials/participants were calculated by summing FCs in the high-anxiety network and summing sign-inversed FCs in the low-anxiety network.

#### 2.1.2. Experimental paradigm

Seven participants attended 6 MRI scanner sessions and 3 participants attended 12 sessions. Each session consisted of 4 blocks, an anxiety block or a neutral block, and each block consisted of 6 trials. The block type order was randomized for all participants. In the first trial of each block, a cue was presented on the screen for 1 s (“anxiety” in the anxiety block, “neutral” in the neutral block). Twenty stimuli were then presented at a rate of 200 ms per stimulus, to minimize the effects of the basic feature and semantic content of a particular stimulus, and was followed by a 14-s imagination period, during which participants were instructed to be anxious if they were in the anxiety block or to relax if they were in the neutral block. After the imagination period, participants were asked to rate their level of anxiety according to a 4-point Likert scale (1, lowest anxiety; 4, highest anxiety). A fixation cross was superimposed on each stimulus, and participants were instructed to maintain fixation on this cross throughout the scanning session. For each session, in two randomly selected trials, participants needed to push the button in response to the change in fixation color on the display to guarantee that participants maintained fixation. These trials were excluded from the subsequent analysis.

#### 2.1.3. fMRI procedure

A 3-T Siemens Trio scanner (Erlangen, Germany) with a 12-channel head coil was used to perform T2*-weighted echo planar imaging (EPI). We acquired 275 scans for each session with a gradient echo EPI sequence. The first 7 scans were discarded to allow for T1 equilibration. The following scanning parameters were used: repetition time (TR), 2000 ms; echo time (TE), 30 ms; flip angle (FA), 80°; field of view (FOV), 192 × 192 mm; matrix, 64 × 64; 33 oblique slices tilted to the anterior commissure–posterior commissure line (Deichmann et al., 2003); and a 3.5-mm slice thickness without gap. T1-weighted anatomical imaging with an MP-RAGE sequence was performed with the following parameters: TR, 2250 ms; TE, 3.06 ms; FA, 9°; FOV, 256 × 256 mm; matrix, 256 × 256; 208 axial slices; and slice thickness, 1 mm without gap.

#### 2.1.4. Preprocessing

We used Statistical Parametric Mapping 8 (SPM8: Wellcome Department of Cognitive Neurology, http://www.fil.ion.ucl.ac.uk/spm/software/) in MATLAB for preprocessing and statistical analyses. The raw functional (EPI) images were initially corrected for slice-timing and realigned to the mean image of that sequence to compensate for head motion. Structural (T1) images were then co-registered to the mean functional image and segmented into three tissue classes in Montreal Neurological Institute (MNI) space. Using the associated parameters, the functional images were then normalized and resampled in a 2 × 2 × 2 mm^3^ grid. Finally, they were smoothed by a Gaussian full-width at half-maximum of 6 mm.

#### 2.1.5. Interregional correlation analysis

For each participant, a pair-wise, interregional FC was evaluated among 268 regions of interest (ROIs) covering the entire brain (Finn et al., 2015). We extracted the representative time course in each region by averaging the time courses of the voxels therein. The time courses were bandpass filtered (transmission range, 0.045–0.18 Hz) and then linearly regressed by the temporal fluctuations in white matter, cerebrospinal fluid, and entire brain, as well as six head motion parameters. We determined the fluctuation in each type of tissue from the average time course of the voxels within a mask created by the segmentation procedure of the T1 image. The mask for the white matter was eroded by one voxel to consider a partial volume effect. These extracted time courses were bandpass filtered (transmission range, 0.045–0.18 Hz) before the linear regression, as was performed for the regional time courses. We defined 10 scans, offset 4 s from the stimulus onset (to account for the delay in hemodynamic response), as the trial scans. For each participant, a matrix of the FCs of each trial between all ROIs was then calculated using scans belonging to each trial by evaluating pair-wise temporal Pearson correlations of BOLD time courses (Fig. 1b). Because FC matrices are symmetric, values on only one side of the diagonal were kept, resulting in 35,778 unique FCs (268 × 267/2) for each trial (Fig. 1c). A final trial in each session has not enough trial scans defined above, that is 10 scans offset 4s from the stimulus onset. Therefore, we excluded the trial from subsequent analysis.

#### 2.1.6. Within-participant anxiety detection

To test the hypothesis that there is a common neural network of anxiety within participants, we first investigated the difference in FCs between trials in different blocks (i.e., the anxiety and neutral blocks) in a fully cross-validated manner (Fig. 2a) for each participant. Cross-validation tests the ability of the model to generalize and involves separating the data into subsets. We analyzed all datasets in a single scale, which was defined by the FCs and named an “anxiety score” (Fig. 1d) (Shen et al., 2017). The score is defined by a subset of the data (the “training” set), and then the generalizability of the score is tested in the fully independent remainder of the data (the “test” set). Here, we conducted 10-fold cross-validation, that is, we split all trials into 10 sets of trials for each participant. For each FC, we then conducted a two-sample *t*-test between the mean values of FC of anxiety and neutral trials using the all but one held-out test set. The resulting *P* values were statistically thresholded at *P* < 0.01 and separated into a positive tail (a higher FC value means that the participant was more likely to see anxiety stimuli) and a negative tail (a higher FC value means that the participant was more likely to see neutral stimuli). We named these FCs high- and low-anxiety networks, respectively. Finally, to validate whether our model reliably detects anxiety in the test set, anxiety scores for trials in the test set were calculated by summing FCs in the high-anxiety network and summing the sign-inversed FCs in the low-anxiety network.

**Figure 2.**
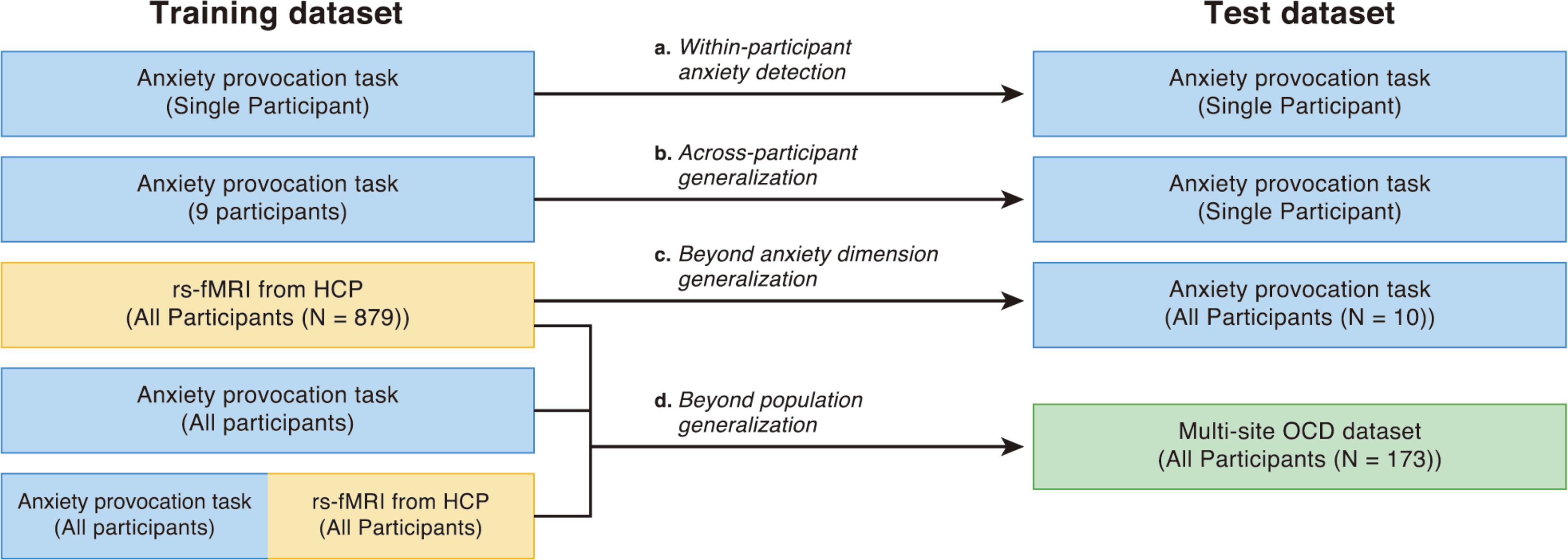
Schematic diagram of the analyses. Anxiety scores were defined by the training dataset and generalized to the held-out test dataset. **(a)** Within-participant anxiety detection was conducted in a 10-fold cross-validation manner. **(b)** Across-participant generalization was conducted in a leave-one-participant-out cross-validation manner. **(c)** Beyond anxiety dimension generalization was conducted using the anxiety score that was defined by the HCP data. **(d)** Beyond population generalization was conducted using anxiety scores defined by the FCs related to state and/or trait anxiety. We also conducted leave-one-participant-out classification within multi-site OCD datasets.

#### 2.1.7. Across-participant generalization

As an additional step, we investigated whether the FCs selected from other participants could be generalized to held-out participants. In other words, we tested whether there was a common neural network of state anxiety across participants. To investigate this, we expanded the previous analysis from the within-participant paradigm to the across-participant paradigm (Fig. 2b). Specifically, for each participant, we calculated the anxiety scores of all trials using almost the same procedure as described in the previous section. However, here, we defined the anxiety score by the other participants’ trials. Thus, we conducted analyses in a leave-one-participant-out cross-validation manner.

### 2.2. Resting-state fMRI: Human Connectome Project

To investigate whether there is a common neural network between state and trait anxiety in healthy people, we tested whether the anxiety score defined by rs-fMRI with the trait anxiety scale can be generalized to state anxiety.

#### 2.2.1. Participants and behavioral measure

We used a public rs-fMRI dataset from the Human Connectome Project (HCP) 900 Subject Release (Van Essen et al., 2012). These data were acquired using a protocol with advanced multiband rs-fMRI sequences for 15 min. The datasets were preprocessed through the common preprocessing pipeline of the HCP (Griffanti et al., 2014; Salimi-Khorshidi et al., 2014). As a behavioral measure of trait anxiety, the NIH Toolbox Fear-Affect Survey, which is a computerized adaptive test comprising items from the PROMIS Anxiety Item Bank (Pilkonis et al., 2011), was used. Participants with both rs-fMRI and behavioral measures were included in the subsequent analyses. The final set of participants comprised 879 participants. More details on these data are described in the Supplementary Notes.

#### 2.2.2. Beyond anxiety dimension generalization

For each participant in the HCP data, using the same ROIs used in the analysis for the state anxiety, the representative time course in each region was extracted by averaging the time courses of the voxels therein. For each participant, a matrix of FC between all ROIs was then calculated using scans by evaluating pair-wise temporal Pearson correlations of BOLD signal time courses of whole scans (Fig. 1b and 1c). We then extracted the FCs related to trait anxiety by calculating Pearson correlations between all FCs and participants’ behavioral measure of trait anxiety (Fig. 1d). Similar to the analyses for the state anxiety alone, the resulting *P* values were statistically thresholded at *P* < 0.01 and separated into a positive tail (a higher FC value means that the participant was more likely to have a high trait anxiety score) and a negative tail (a higher FC value means that the participant was more likely to have a low trait anxiety score). Again, we called these FCs high- and low-anxiety networks, respectively. The anxiety score was calculated for all participants’ trials in the anxiety provocation task, as previously described (Fig. 2c).

### 2.3. Common neural network between state and trait anxiety

To evaluate the degree of overlap between sets of FCs related to state and trait anxiety, we then tested whether the number of common FCs was significantly higher than chance. The FC was considered to be common if it was included in the same network (high- or low-anxiety network) of both state and trait anxiety. Note that the FCs related to state anxiety were defined by FCs that were selected at all iterations through the leave-one-participant-out cross-validation procedure for the anxiety provocation task. The FCs related to trait anxiety were defined by FCs that were selected using the HCP data. To determine the number of common FCs obtained by chance, we compared randomly created networks of state and trait anxiety. First, we shuffled the feature matrix of state anxiety and defined a high- and low-anxiety network as described in the previous section. We then also shuffled the feature matrix of trait anxiety. We repeated this procedure 10,000 times and defined 10,000 random high- and low-anxiety networks. Finally, we calculated the number of common FCs between state and trait anxiety for every random low- and high-anxiety network.

### 2.4. Prediction of patients with OCD

Even if there is a set of FCs related to state and/or trait anxiety in healthy people, the relationships between these networks and pathological anxiety is still a question that remains to be directly investigated. Therefore, we asked which set of FCs is altered in the population with pathological anxiety among the FCs related to state and/or trait anxiety. To investigate this question, we used three rs-fMRI datasets consisting of two different populations, namely, healthy controls (HCs) and patients with OCD, which is characterized by pathological anxiety-driven behavior.

#### 2.4.1. Participants

We recruited participants at three different sites: sites A and B in Japan and site C in Spain. Supplementary Table 1 shows a summary of the participants’ demographic information. Each imaging site adopted its own imaging protocol (Supplementary Table 2), differing in imaging parameters. The complete recruitment criteria are described in the Supplementary Notes.

**Table 1:**
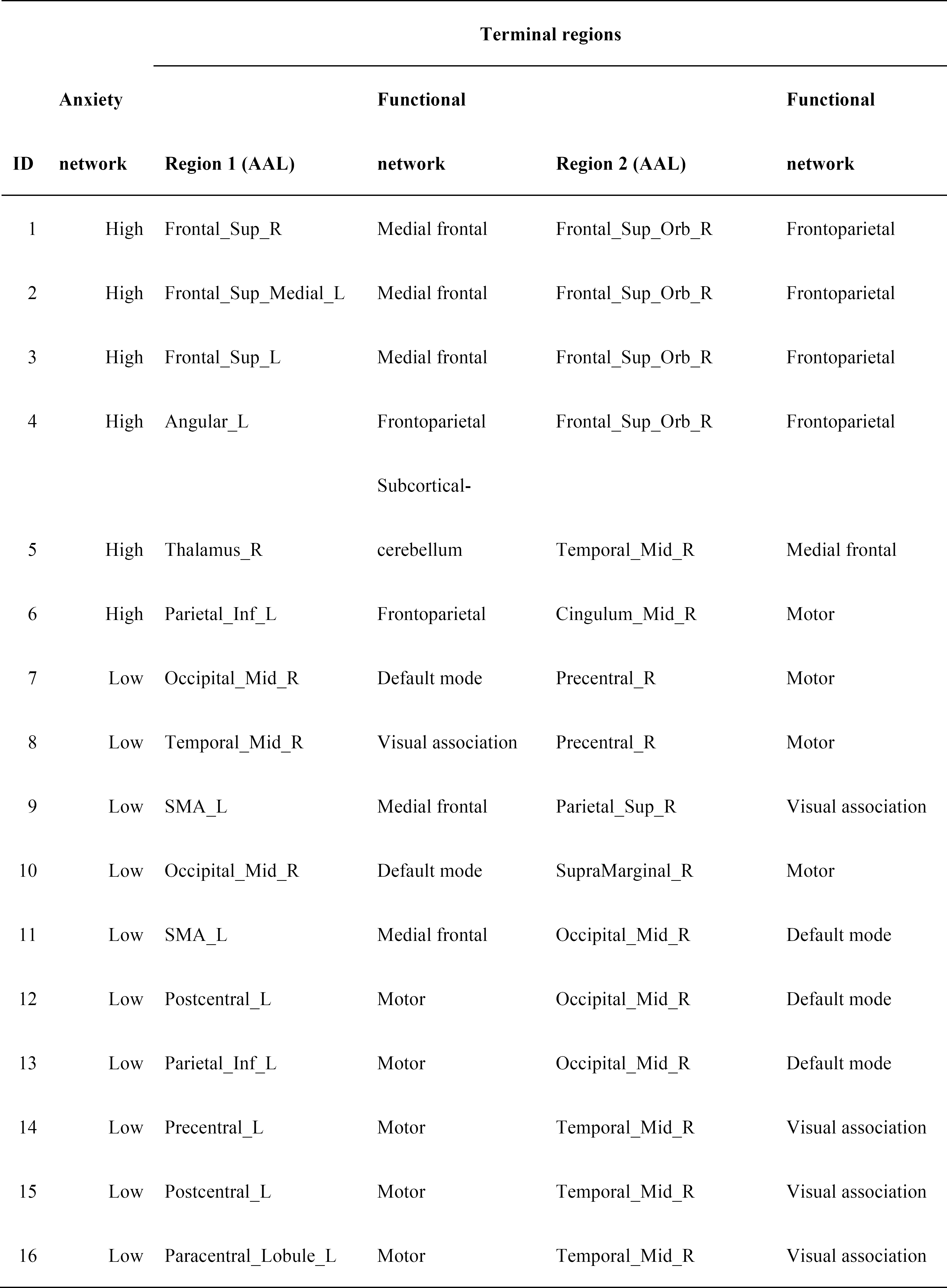
Properties of the 16 common FCs between state and trait anxiety. Mid, Middle; Sup, Superior; Inf, Inferior; Orb, Orbitofrontal; SMA, Supplemental Motor Area; R, Right; L, Left.

At site A (Kajiicho Medical Imaging Center, Kyoto, Japan), the same dataset used by Takagi et al. (2017) was used. All resting-state fMRI data (56 OCD and 52 HCs) were collected at Kajiicho Medical Imaging Center, Kyoto, Japan. The Medical Committee on Human Studies at Kyoto Prefectural University of Medicine, Kyoto, Japan, approved all of the procedures in this study. All participants gave written informed consent after receiving a complete description of the study. For each participant, a single 6-min 40-s continuous resting functional scan and a high-resolution T1-weighted anatomical image were acquired.

At site B (KPUM), the same dataset used by Sakai et al. (2011) was used. Because 15 participants were also included in site A, we excluded them from site B. Finally, 28 participants were included (10 OCD and 18 HCs). The Medical Committee on Human Studies at Kyoto Prefectural University of Medicine, Kyoto, Japan, approved all of the procedures in this study. All participants gave written informed consent after receiving a complete description of the study. For each participant, a single 8-min continuous resting functional scan and a high-resolution T1-weighted anatomical image were acquired.

At site C (Bellvitge University Hospital, Barcelona, Spain), the same dataset reported by Harrison et al. (2009) was included. Forty-two participants were recruited from the Department of Psychiatry, Bellvitge University Hospital, Barcelona, Spain. Because data were not available on 4 of the 21 HCs, we excluded them from our analysis. Finally, 21 patients with OCD and 17 HCs were included in the dataset. All participants gave written informed consent after receiving a complete description of the study. The institutional review board of the Bellvitge University Hospital, Barcelona, approved all of the procedures in this study. For each participant, a single 4-min continuous resting functional scan and a high-resolution T1-weighted anatomical image were acquired.

#### 2.4.2. Preprocessing

We used SPM8 for preprocessing and statistical analyses in a similar manner to our previous rs-fMRI studies (Takagi et al., 2017). Raw functional images were first corrected for slice-timing and realigned to the mean image of that sequence to compensate for head motion. Structural images were then co-registered to the mean functional image and segmented into three tissue classes in the MNI space. Using associated parameters, the functional images were then normalized and resampled in a 2 × 2 × 2 mm^3^ grid. Finally, they were smoothed by a Gaussian full-width at half-maximum of 6 mm. To avoid the effects of motion artifacts, we employed the “scrubbing” procedure to identify and exclude any frames exhibiting excessive head motions (Power et al., 2012). A bandpass filter (transmission range, 0.008–0.1 Hz) was then applied to the sets of time courses. The filtered time courses were linearly regressed by the temporal fluctuations of the white matter, cerebrospinal fluid, and entire brain, as well as six head motion parameters. These extracted time courses were bandpass filtered (0.008–0.1 Hz) before the linear regression, as was performed for regional time courses. Finally, for each participant, using the same ROIs and procedure for the rs-fMRI from the HCP data, the rs-FC matrix was calculated (Fig. 1b and 1c). The complete procedure is described in the Supplementary Notes.

#### 2.4.3. Beyond population generalization

To investigate how state and trait anxiety are involved in pathological anxiety, we tested which sets of FCs were generalized to patients with OCD, among the neural networks related to state and/or trait anxiety. That is, we calculated three types of the anxiety score for each participant in the multi-site OCD datasets. Their anxiety scores were calculated by summing the FCs in the high-anxiety network and summing the sign-inversed FCs in the low-anxiety network (Fig. 1d). These networks were defined by state and/or trait anxiety data (Fig. 2d). We then compared the anxiety scores of HCs with those of patients with OCD or state and/or trait anxiety.

#### 2.4.4. Machine-learning classification

Recently, rs-fMRI has received attention as a biomarker of psychiatric disorder in the clinical field (Fox and Greicius, 2010). Therefore, we tested whether the selected high- and low-anxiety networks could also be used as a biomarker of OCD. Note that, although several studies have investigated neurological biomarkers of OCD (Gruner et al., 2014; Hu et al., 2016; Li et al., 2014; Soriano-Mas et al., 2007; Takagi et al., 2017; Weygandt et al., 2012), our study is distinct because we did not use any diagnostic label and focused on the dimensions of the anxiety. We conducted the classification via leave-one-participant-out cross-validation employing a sparse logistic regression classifier (Yamashita et al., 2008). Thus, the weights of the classifier were estimated using training participants and were validated using test participants. For this analysis, we used the common FCs between state and trait anxiety because only this set of FCs was generalized to pathological anxiety in the previous analysis (see the Results). Furthermore, to demonstrate that the results cannot be reproduced by randomly selected FCs, we picked the same number of FCs randomly from all 35,778 FCs and then conducted the same procedure 10,000 times to determine the distribution of the area under the curve (AUC).

### 2.5. Interpretation of anxiety-related FC patterns

To facilitate characterization of the biological substrates of the anxiety-related neural network, we summarized the FC patterns that were obtained from the state anxiety, trait anxiety, and their overlap. We grouped the 268 ROIs into eight macroscale canonical networks (e.g., the default mode network) and examined the number of FCs between each pair of regions in each network; these canonical networks were defined functionally in a previous study (Finn et al., 2015).

## 3. Results

### 3.1. State anxiety-related network: anxiety provocation task

#### 3.1.1. Within-participant anxiety detection

To test whether there is a neural network that is consistently different between neutral and anxious states within participants, we compared the anxiety scores of the trials in different blocks in a cross-validated manner for each participant (Fig. 2a). We tested whether the anxiety scores of trials in the neutral block were lower than those of trials in the anxiety block. We applied a two-way repeated measures analysis of variance (ANOVA) to the anxiety scores of all trials (N = 1638; participant × stimulus [anxiety or neutral]) and identified a significant effect of stimulus (F_(1,1637)_ = 209.66; *P* = 9.03 × 10^−45^). Figure 3a shows the anxiety scores of the anxiety and neutral trials, which were normalized within each participant. We also found a significant effect of participant (F_(9,1637)_ = 88.43; *P* = 7.53 × 10-134) and interaction (F_(9,1637)_ = 17.27; *P* = 1.64 × 10^−27^). Note that the anxiety scores of all trials were defined from other trials (i.e., we did not use them themselves in order to avoid circularity).

**Figure 3.**
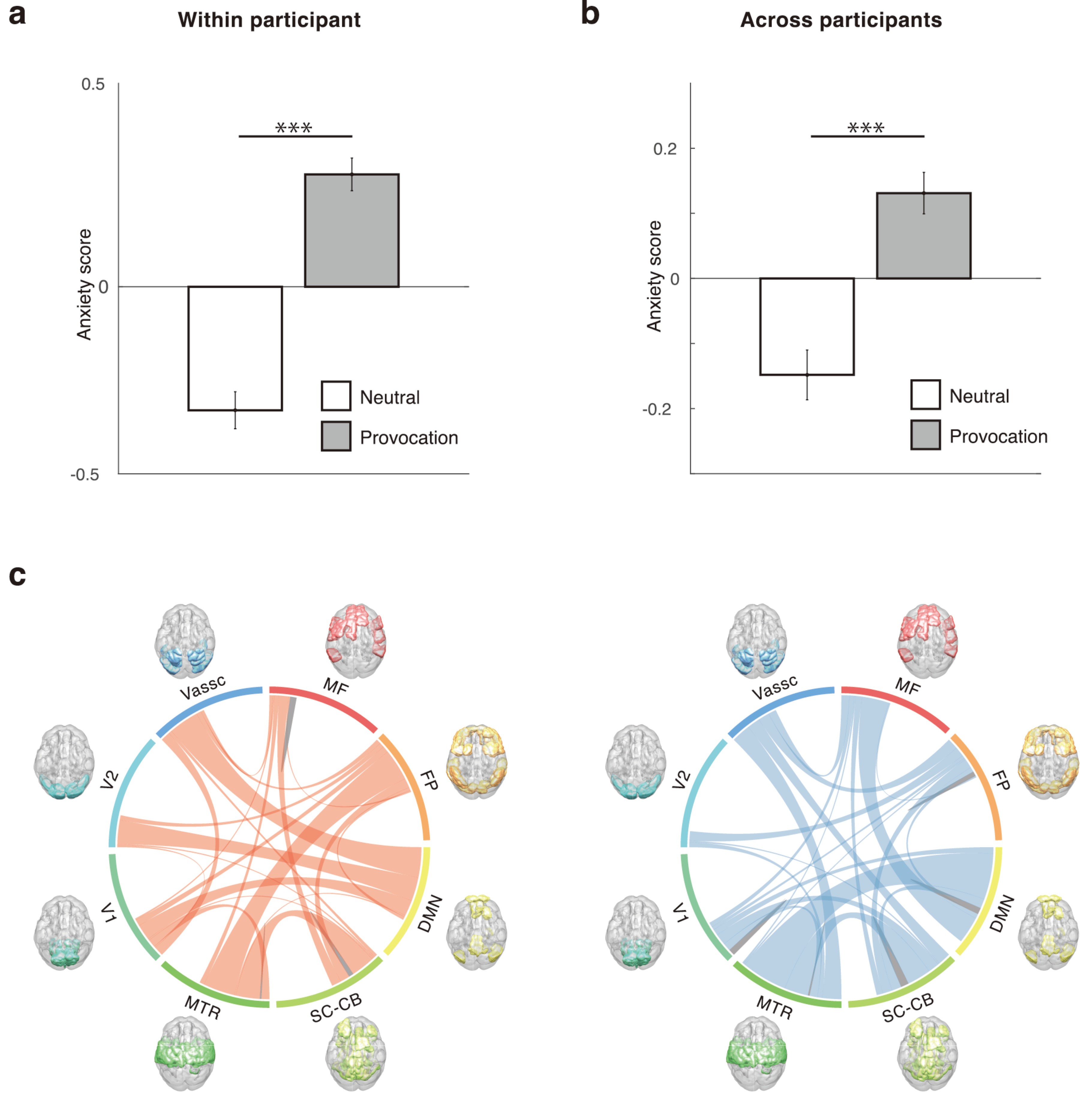
State anxiety-related neural network defined by the anxiety provocation task. **(a–b)** The anxiety scores of the anxiety stimuli (gray) and neutral stimuli (white) for **(a)** within- and **(b)** across-participant analyses. For visualization purposes, the anxiety scores of each participant were normalized within each participant. Note that this normalization was not performed in any quantitative analysis. Error bars indicate SEM. **(c)** Macroscale visualization of high-anxiety (left, red) and low-anxiety (right, blue) networks for the number of FCs that consistently differed between the trials of anxiety stimuli and neutral stimuli across participants. Canonical networks include the medial frontal (MF), frontoparietal (FP), default mode network (DMN), subcortical-cerebellum (SC-CB), motor (MTR), visual I (V1), visual II (V2), and visual association (VAssc). Connection lines are colored gray within the same network and red (high) or blue (low) between two networks. ^***^Two-way repeated measures ANOVA, *P* < 0.001.

#### 3.1.2. Across-participant generalization

To investigate whether the neural network is different between neutral and anxious states in the same manner across participants, we expanded the previous analysis from the within-participant paradigm to the across-participant paradigm (Fig. 2b). We applied two-way repeated measures ANOVA to the anxiety scores of all trials (N = 1638; participant × stimulus [anxiety or neutral]) and obtained similar results to the previous analysis, that is, the effect of the stimulus was significant (F_(1,1637)_ = 24.26; *P* = 9.29 × 10^−7^). Figure 3b shows the anxiety scores of the trials of anxiety stimuli and neutral stimuli, which were normalized within participants. We also found a significant effect of participant (F_(9,1637)_ = 18.87; *P* = 3.05 × 10^−30^) but no interaction effect (F_(9,1637)_ = 1.36; *P* = 0.20).

#### 3.1.3. Visualization of the state anxiety-related neural network

Through the leave-one-participant-out cross-validation procedure, we observed hundreds of FCs selected at all iterations, that is, hundreds of FCs consistently differed between the trials of anxiety stimuli and neutral stimuli across participants. To facilitate characterization of the biological substrates of the state anxiety-related FCs, we grouped the 268 ROIs into eight macroscale canonical networks. Figure 3c shows the circle plot of the FCs that were involved in the high-anxiety network and the low-anxiety network. The numbers of FCs in each of the two macroscale regions (the medial frontal [MF], frontoparietal [FP], default mode network [DMN], subcortical-cerebellum [SC-CB], motor [MTR], visual I [V1], visual II [V2], and visual association [VAssc]) are presented as the thickness of the connection lines. Although the FCs in the DMN were selected to some extent, the FCs in both networks were widely distributed rather than locally constrained.

### 3.2. Beyond anxiety dimension generalization: resting-state fMRI from the Human Connectome Project

To investigate whether there is a common neural network between state and trait anxiety, we examined all trials of the anxiety provocation task (N = 1638) using the anxiety score that was defined by the HCP data with the rs-fMRI and trait anxiety scale (N = 879) (Fig. 2c). We applied a two-way repeated measures ANOVA to the anxiety scores of all trials (participant × stimulus [anxiety or neutral]) and the results showed that the anxiety scores of the trials of anxiety stimuli and neutral stimuli were significantly different (F_(1,1637)_ = 7.72; *P* = 5.54 × 10^−3^). Figure 4a shows the normalized anxiety scores within participants. We also found a significant effect of participant (F_(9,1637)_ = 35.86; *P* = 3.12 × 10^−58^) and interaction (F_(9,1637)_ = 3.2; *P* =7.64 × 10^−4^). Figure 4b shows the high- and low-anxiety networks in a macroscale defined by the HCP data with the rs-fMRI and trait anxiety scale. The high- and low-anxiety networks of trait anxiety revealed more dissimilar patterns of the FCs than those of state anxiety. In particular, the FCs within the frontoparietal and medial-frontal networks and between them were frequently selected in the high-anxiety network.

**Figure 4.**
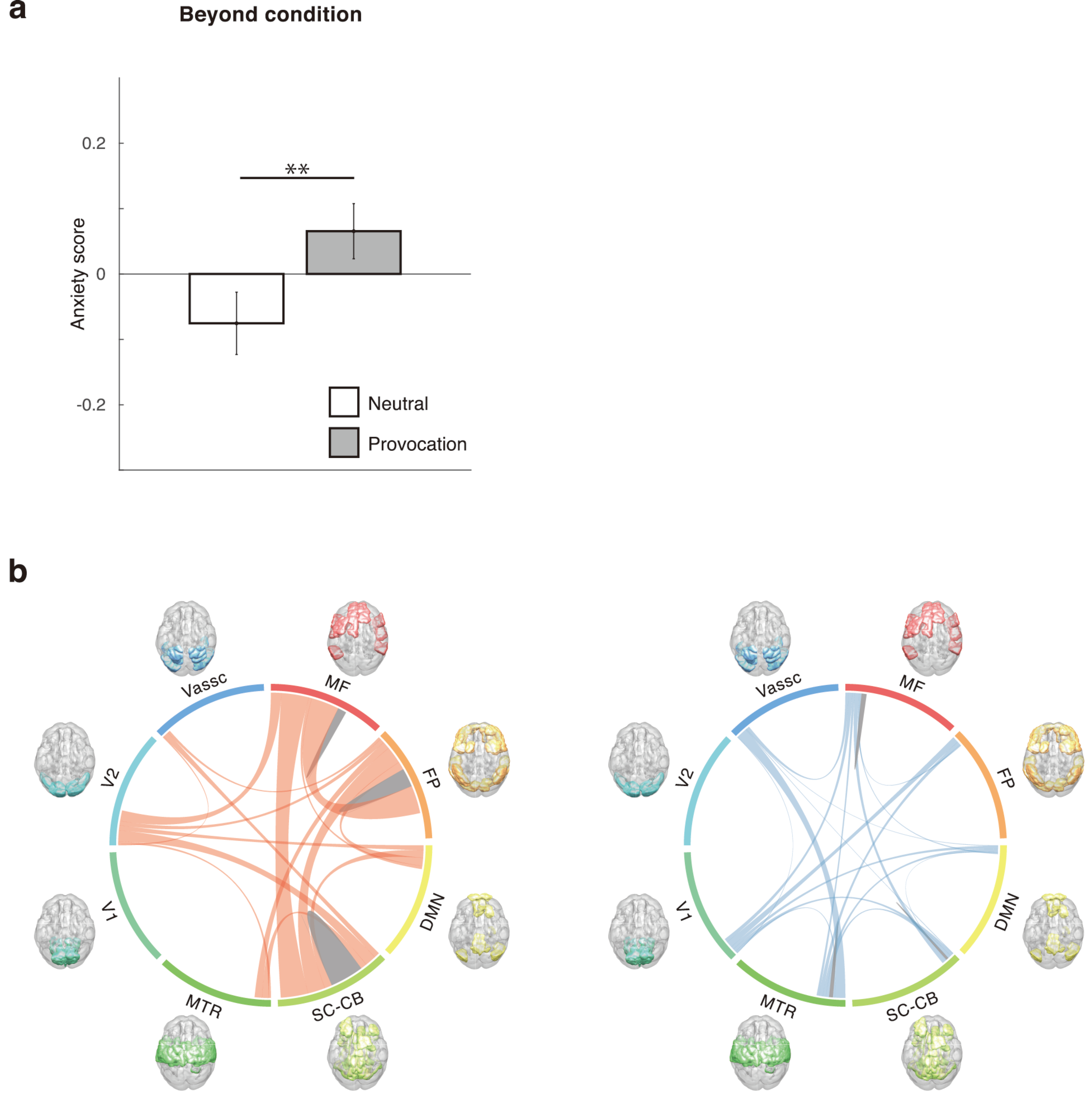
Generalization to different dimensions of anxiety. **(a)** The anxiety scores of the anxiety stimuli (gray) and neutral stimuli (white). Error bars indicate SEM. **(b)** High-anxiety (left, red) and low-anxiety (right, blue) networks defined by the HCP data with the rs-fMRI and trait anxiety scale in a macroscale. The numbers of FCs in each of the two macroscale regions are presented as the thickness of the connection lines. Connection lines are colored gray within the same network and red (high) or blue (low) between two networks. Abbreviations are the same as in Figure 3. ^**^Two-way repeated measures ANOVA, *P* < 0.01.

### 3.3. Common neural network between state and trait anxiety

To evaluate the degree of overlap between neural networks related to state and trait anxiety, we counted the common FCs between these two networks. Among the whole 35,778 FCs used, 16 FCs were included in the same network (the high- or low-anxiety network) in both dimensions of anxiety (state or trait anxiety) (Table 1). Specifically, when we extracted FCs using the whole dataset of state anxiety, 1,187 FCs were selected, comprising 573 FCs in the high-anxiety network and 614 FCs in the low-anxiety network. For trait anxiety, 382 FCs were selected, comprising 287 FCs in the high-anxiety network and 95 FCs in the low-anxiety network. Finally, 6 and 10 FCs overlapped in the high- and low-anxiety networks, respectively. Figure 5a shows the spatial distribution of these 16 FCs. Four of the 6 FCs in the high-anxiety network had the orbitofrontal cortex as their node. The other 2 FCs had the thalamus and cingulate cortex as their nodes. In contrast, the FCs of the low-anxiety network were in the motor cortex and occipital cortex. For macroscopic interpretation, Fig. 5b shows the common FCs in a macroscale. The FCs in the high-anxiety network frequently belonged to the frontoparietal and medial-frontal network. The FCs in the low-anxiety network belonged to the default mode network, subcortical-cerebellum, and motor network. Finally, we compared 10,000 randomly generated anxiety networks with our results and found that the probability for obtaining the 16 overlapped FCs was statistically significant (*P* < 0.001). This suggests that the 16 FCs were not merely randomly overlapping but play a significant 4 role as a common neural network of anxiety.

**Figure 5.**
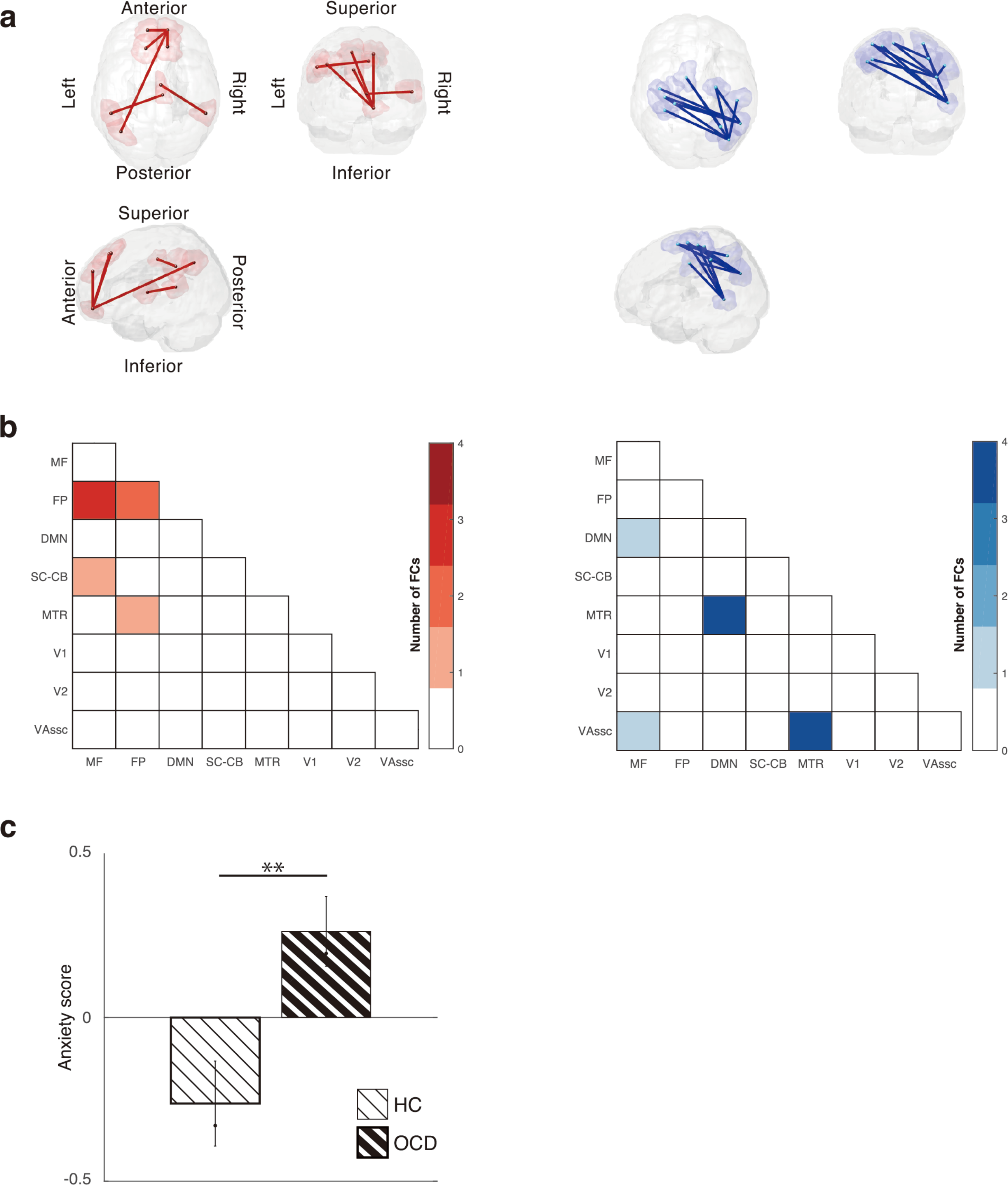
The common neural network and its generalization to patients with obsessive-compulsive disorder at the three different sites. **(a)** Spatial distribution of the 16 common FCs. **(b)** Matrices for the number of FCs between each pair of canonical networks in the high-anxiety (left, red) and low-anxiety (right, blue) networks. Abbreviations are the same as in Figure 3. **(c)** The anxiety scores of the healthy controls (white) and patients with OCD (gray). Error bars indicate SEM. ^**^Two-way repeated measures ANOVA, *P* < 0.01.

### 3.4. The common neural network of anxiety among different populations: patients with OCD from three different sites

To investigate which neural network is involved in pathological anxiety, we compared the anxiety scores defined by state anxiety and/or trait anxiety of patients with OCD and HCs from the three different datasets (N = 174) (Fig 2d). We applied two-way repeated measures ANOVA to the anxiety scores of all trials (population [HC or OCD] × sites [site A, B, or C]). We found that only when we applied the anxiety score defined by the 16 common FCs, the anxiety scores of patients with OCD were significantly different from those of HCs (Fig. 5c; F_(1,173)_ = 8.56; *P* = 3.91 × 10^−3^). We also found a significant effect of site (F_(2,173)_ = 4.8; *P* = 9.39 × 10^−3^) but no interaction effect (F_(2,173)_ = 0.46; *P* = 0.63). Notably, when we conducted the same analyses using the anxiety scores defined by either state or trait anxiety, no significant differences were observed for any comparisons (*P* > 0.05 for all comparisons).

### 3.5. The common neural network as a biomarker: classification of patients with OCD

Finally, to test whether the 16 common FCs specified above could also be used as a biomarker of OCD, we evaluated the performance of a classifier that was composed of these FCs. Our model, developed by the 16 common FCs, achieved a significantly higher AUC (AUC = 0.63) than other models that consisted of the same number of randomly selected FCs (1,000 times permutation test; *P* < 0.02). This analysis indicates that the probability of obtaining the AUC was small and demonstrates that the 16 FCs identified in the main analyses specifically contain information useful to predict OCD.

## 4. Discussion

In this study, we have characterized a common neural network among state, trait, and pathological anxiety by analyzing different fMRI datasets that were bound to different dimensions of anxiety in a data-driven manner. We first demonstrated that there was a common neural network of state anxiety within and between participants. We then demonstrated that the trait anxiety-related neural network, defined by rs-fMRI with the trait anxiety scale, can be generalized to state anxiety. Finally, we found that we could characterize pathological anxiety, as represented by patients with OCD, by using the common FCs between state and trait anxiety.

Although previous fMRI studies have reported patterns of neural activation during anxiety provocation tasks (Blair et al., 2008; Lorberbaum et al., 2004; Mataix-Cols et al., 2003; Nakao et al., 2005; Tillfors et al., 2001), they did not investigate whole-brain FCs and did not perform cross-validation. In the present study, we first demonstrated that the common FCs related to state anxiety across participants can be detected in a fully cross-validated manner. These results suggest that when we feel anxiety in our daily life, common brain regions might be co-activated within and between individuals.

Our findings also revealed that there is a common neural network between state and trait anxiety. Our results suggest that the resting-state FC patterns of participants who tend to feel anxiety in their daily life is similar to the FC pattern during evoked anxiety. This finding is noteworthy because previous studies showed that state and trait anxiety often interact (Mathews, 1990; Williams et al., 1996). In these studies, the authors hypothesized that participants with high trait anxiety become increasingly vigilant under stress, further increasing their anxiety level. In contrast, participants with low trait anxiety show a defensive response under stress, serving to restrain further anxiety increases. Common FCs may contribute to this positive-negative feedback loop of anxiety.

In the present study, we identified common FCs between state and trait anxiety, and the number of common FCs was significantly larger than in randomly selected cases. Furthermore, although we employed a fully data-driven approach rather than setting an a priori hypothesis, the pattern of FCs revealed was highly meaningful. That is, frontal and default mode networks were more likely chosen, and they were often reported in previous studies as anxiety-related regions (Beucke et al., 2013; Liao et al., 2010; Liu et al., 2015; Modi et al., 2015; Tian et al., 2016; Wang et al., 2016; Yin et al., 2016). More specifically, the orbitofrontal cortex, thalamus, and cingulate cortex were included in the high-anxiety network and the default mode network was included in the low-anxiety network. The increased and decreased FCs in these areas for state, trait, and pathological anxiety have frequently been reported. It should be noted that, as shown in several seed-based studies, the amygdala has often been reported as a central region of anxiety (Banca et al., 2015; Baur et al., 2013; Cha et al., 2014; Hahn et al., 2011; Kim et al., 2011). In the present study, we found no evidence indicating that the amygdala is fundamentally involved in anxiety, as operationalized here. Intriguingly, the above studies using data-driven procedures commonly do not report the amygdala and, additionally, most have reported a frontal and/or default mode network. To our knowledge, no data-driven study of anxiety has investigated the generalizable FCs among different dimensions of anxiety. Our study has quantitatively evaluated the relative importance of the conventionally investigated brain regions to whole brain using a fully data-driven approach.

When we used the common FCs between state and trait anxiety to calculate the anxiety scores of patients with OCD and HCs, patients with OCD showed higher anxiety scores than HCs. Furthermore, the FCs could classify whole participants into HCs and patients with OCD in a fully cross-validated manner. However, we could not find such results when we used anxiety scores defined by either state or trait anxiety data. Our findings thus demonstrate that the common FCs between state and trait anxiety are key components of pathological anxiety. Our findings may also be theoretically helpful for constructing a biologically validated model of anxiety that explicitly considers the interaction among state, trait, and pathological anxiety. In addition, future work should also investigate the causal interaction among state, trait, and pathological anxiety by modulating the neural network, for example, via neurofeedback (Megumi et al., 2015), to obtain a deeper understanding of the neural mechanism of anxiety. Our findings may contribute to the identification of a reasonable target for intervention.

## 5. Conclusion

In this study, we found a common neural network of anxiety among state, trait, and pathological anxiety. These results provide direct evidence that different dimensions of anxiety are not completely independent but have a substantial biological inter-relationship. Our results also have the potential to be used for the prevention of pathological anxiety before onset or for treatment evaluation. Furthermore, given that the hypothesis of a psychiatric disorder spectrum is gaining attention, our results may promote further investigation of various human characteristics, including psychiatric disorder, from the perspective of anxiety.

## Acknowledgementss

This research was conducted as the “Application of DecNef for development of diagnostic and cure system for mental disorders and construction of clinical application bases” of the Strategic Research Program for Brain Sciences from the Japan Agency for Medical Research and Development, AMED. This work was partly supported by JSPS KAKENHI Grant Number 25119001, the Joint Usage/Research Center at ISER, Osaka University, and a grant from the Carlos III Health Institute (PI13/01958). Dr. Soriano-Mas is funded by a Miguel Servet contract from the Carlos III Health Institute (CPII16/00048). Data were provided [in part] by the Human Connectome Project, WU-Minn Consortium (Principal Investigators: David Van Essen and Kamil Ugurbil; 1U54MH091657) funded by the 16 NIH Institutes and Centers that support the NIH Blueprint for Neuroscience Research; and by the McDonnell Center for Systems Neuroscience at Washington University. We thank Okito Yamashita and Hiroshi Imamizu for helpful discussions and proofreading the manuscript. We thank T. Okada, H. Ito, and the technical engineers for their assistance in MRI data acquisition. We thank K. Tamura, S. Kimura, and K. Inoue for their assistance in the assessment of patients. We thank N. Izumihara for his support for visualization.

